# Phenotypic characterization and analysis of complete genomes of two distinct strains of the proposed species “L. swaminathanii”

**DOI:** 10.1101/2022.04.22.489179

**Authors:** Lauren K. Hudson, Harleen K. Chaggar, Claire N. Schamp, Michelle L. Claxton, Daniel W. Bryan, Tracey L. Peters, Yaxiong Song, Catharine R. Carlin, Henk C. den Bakker, Thomas G. Denes

## Abstract

Recently, a new *Listeria* species, “Listeria swaminathanii”, was proposed. Here, we phenotypically and genotypically characterize two additional strains that were previously obtained from soil samples and compare the results to the type strain. Complete genomes for both strains were assembled from hybrid Illumina and Nanopore sequencing reads and annotated. Further genomic analysis including average nucleotide identity (ANI) and detection of mobile genetic elements and genes of interest (e.g., virulence-associated) were conducted. The strains showed 98.7-98.8% ANI with the type strain. The UTK C1-0015 genome contained a partial monocin locus and a plasmid while the UTK C1-0024 genome contained a full monocin locus and a prophage. Phenotypic characterization consistent with those performed on the proposed type strain was conducted to assess consistency of phenotypes across a greater diversity of the proposed species (n=3 instead of n=1). Only a few findings were notably different from those of the type strain, such as catalase activity, glycerol metabolism, starch metabolism, and growth at 41°C. This study further expands our understanding of this newly proposed *sensu stricto Listeria* species.

## INTRODUCTION

*Listeria* spp. are small Gram-positive, motile, non-sporulating, and non-capsulated rods^1–3^. The *Listeria* genus consists of two clades, *sensu stricto* and *sensu lato*^4,5^, and currently contains 26 validly published species^6^. The *sensu stricto* clade includes the human and animal pathogen *L. monocytogenes*^7^. *Listeria* spp. are ubiquitous and commonly isolated from natural environments^8^ and the majority of novel species or subspecies described in recent years were originally isolated from these environments^4,9–15^.

Recently, a strain isolated from soil collected in the Great Smoky Mountains National Park (GSMNP)^8^ was proposed as a novel *sensu stricto* species, “Listeria swaminathanii”^16^. However, it was unable to be validly published due to culture collection deposition restrictions imposed by the National Park Service and rules on strain availability set forth by the International Committee on Systematics of Prokaryotes (ICSP).

In another study, Claxton and Hudson et al.^17^ obtained two distinct isolates, UTK C1-0015 and UTK C1-0024, from soil samples collected in the GSMNP that were not closely related to any published type species. Here, we show that these two isolates are additional members of the proposed species “L. swaminathanii”. We further characterized these two strains genotypically and phenotypically to expand our understanding of this newly proposed species by increasing the number of characterized strains from one to three. Additionally, we sequenced the isolates using both short- and long-read sequencing technologies and were able to produce complete closed genomes and further characterized the genomic features of each isolate.

## MATERIALS & METHODS

### Genome sequencing and assembly

Genomic DNA was extracted using a Qiagen QIAamp DNA mini kit (Hilden, Germany) per manufacturer protocol, with the addition of an RNase treatment step^18^. For short-read sequencing, library preparation and sequencing were performed by the Microbial Genome Sequencing Center (MiGS; Pittsburgh, PA). Sequencing was performed on an Illumina NextSeq 2000 instrument with 151 bp paired-end read chemistry. For each, 2.5-2.6 million total paired sequencing reads were produced, with average lengths of 146.0-146.3 bp. Mean quality phred scores were >32, indicating good quality calls.

For long-read sequencing, the SQK-RBK004 kit (Oxford Nanopore Technologies, Oxford, UK) was used for library preparation and a MinIon instrument with a FLO-MIN106 flow cell were used for sequencing, along with the MinKNOW software (v3.6.5, fast basecalling model). A total of 293,459 total sequencing reads were produced for UTK C1-0015 with an average length of 5,656 bp. A total of 86,653 sequencing reads were produced for UTK C1-0024 with an average length of 5,862 bp.

Raw Illumina reads were trimmed using Trimmomatic^19^ (v0.39; with the following parameters: ILLUMINACLIP:NexteraPE-PE.fa:2:30:10 LEADING:3 TRAILING:3 SLIDINGWINDOW:4:15 MINLEN:36). Read quality statistics for both types of reads were generated with FastQC^20^ (v0.11.9). Both short- and long-reads were used to create hybrid assemblies using Unicycler^21^ (v0.4.8; default parameters). Assembly statistics were generated with QUAST^22^ (v5.0.2), BBMAP^23^, and SAMtools^24^ (v1.10). Assemblies were submitted to NCBI and annotated using the NCBI Prokaryotic Genome Annotation Pipeline^25^ (PGAP; v5.3).

### Genomic Characterization

Relatedness to other species and taxonomy were assessed using various genomic methods, including ANI, ribosomal multilocus sequence typing (rMLST), and dDDH. Assemblies for all currently described *Listeria* species type strains and representative strains were downloaded from the RefSeq or GenBank databases on NCBI or the ATCC genome portal, along with the assembly for the “L. swaminathanii” type strain (<u>GCF_014229645.1</u>). PYANI^26^ (v0.2.10) was used to calculate ANI between all strains and bactaxR^27^ was used to create an ANI dendrogram. The assemblies were also input into the rMLST^28^ tool (available on PubMLST) and the Type Strain Genome Server (TYGS)^29^. A whole-genome alignment was performed with the two strains and FSL L7-0020 in Geneious using the progressiveMauve^30^ algorithm (Mauve plugin v1.1.3) and visualized with Mauve^30,31^. For the alignment, the FSL L7-0020 assembly contigs reordered relative to the strain genomes and concatenated into a single sequence to form a pseudochromosome using the MCM algorithm in Geneious; the plasmid was also excluded from UTK C1-0015.

Genomes were evaluated for loci associated with antimicrobial resistance, virulence, motility, metal and disinfectants resistance, stress islands, and *Listeria* genomic islands using ResFinder^32^ (v4.1), KmerResistance^33,34^ (v2.2), VirulenceFinder^35^ (v2.0), and the relevant schemes on Pasteur^36–39^. Mobile genetic elements (MGEs) were identified and characterized using PlasmidFinder (v.2.0), PLSDB^40^ (v.2021_06_23), Phaster^41^, and PhageBoost^42^. BLAST and BLAST Ring Image Generator (BRIG) (v0.95)^43^ were used to create a plasmid map for comparison of similar plasmids. Genomic comparison of monocin loci and nucleotide and amino acid identity were determined using BLAST and EasyFig^44^ (v2.2.2).

#### Phenotypic characterization

The phenotypic characterization of UTK C1-0015 and UTK C1-0024 was performed as per the standardized methodology in the FDA Bacteriological Analytical Manual (BAM) Chapter 10^45^ and those described by Carlin, et al.^10,16^. The following characteristics were assessed: growth at different temperatures (4, 7, 22, 30, 37, and 41°C), growth under anaerobic conditions, colony morphology on selective and differential agar medium, motility, Gram stain, hemolysis, oxidase and catalase activity, nitrate reduction, and the biochemical tests included in three different commercial test kits (API *Listeria*, API 20 E, and API 50 CH). For each phenotypic analysis, from frozen stock, strains were streaked for isolation onto Brain Heart Infusion (BHI) agar (BD Biosciences, Franklin Lanes, NJ+ Fisher Scientific Agar, Waltham, MA) and incubated aerobically at 30°C for 24 h. From that plate, isolated colonies were either used directly or to inoculate a BHI broth (BD Biosciences, Franklin Lanes, NJ) tube, followed by aerobic incubation with shaking at 30°C for 24h. Unless otherwise specified, three biological replicates were performed for each test, each starting from a different single, isolated colony. Control strains included the well-characterized *L. monocytogenes* 10403S, *L. monocytogenes* ATCC 19115, *L. ivanovii* subsp. *ivanovii* ATCC 19119, *L. innocua* ATCC 33090, *L. seeligeri* ATCC 35967, and *L. booriae* FSL A5-0281^T^.

### Growth temperature

To measure growth at different temperatures, BHI broth cultures were used to inoculate 5 mL BHI broth tubes to a concentration of 10^2^-10^3^ CFU/mL for each strain and temperature combination; *L. monocytogenes* 10403S was included as a positive control. The tubes were then aerobically incubated at 4, 7, 22, 30, 37, or 41°C for up to 5 d. Enumerations were performed by spread plating onto BHI agar in duplicate at 24 and 48 h for 22, 30, 37, and 41°C, at 11 and 14 d for 4°C, and at 11 and 15 d for 7°C. Enumeration plates were incubated for 24-36 h at 30°C. If no growth occurred at 48 h, additional enumerations were performed daily for up to 5 d.

### Anaerobic growth

To assess growth under anaerobic conditions, BHI agar plates were streaked from BHI broth cultures in duplicate; *L. monocytogenes* 10403S was included as a positive control. Plates were incubated at 30°C, one aerobically and one anaerobically (in an anaerobic chamber with GasPak™ EZ Anaerobe Container System Sachets with Indicator [BD Difco, Franklin Lanes, NJ]). Growth was assessed at 24 and 48 h.

### Selective and differential agars

For colony phenotypes on selective and differential agars, BHI broth cultures were streaked onto modified oxford agar (MOX; Remel Oxford Agar Base Modified, Lenexa, KS; BD Difco Supplement, Franklin Lanes, NJ) and *Listeria* CHROMagar (commercially prepared; BD Biosciences, Franklin Lanes, NJ) plates. *Listeria monocytogenes* 10403S, *Listeria ivanovii* ATCC 19119, *L. seeligeri* ATCC 35967, and *Listeria innocua* ATCC 33090 were included as controls. Plates were incubated aerobically at 30°C and evaluated at 24 and 48 h.

### Motility

Two methods were used to detect motility: microscopic observation and observation of growth in Motility Test Medium (MTM). *L. monocytogenes* 10403S and *L. booriae* FSL A5-0281^T^ were included as positive and negative controls, respectively. BHI agar streak plates were incubated aerobically at 25°C and 37°C for 24 h. Wet mounts from both sets of plates were observed microscopically for tumbling motility. This assay was completed once for each strain and temperature combination. Additionally, MTM tubes (commercially prepared; Hardy Diagnostics, Santa Maria, CA) were stab inoculated from the 25°C BHI agar plates, incubated at 25°C, and observed daily for 7 d.

### Oxidase and catalase

Oxidase and catalase tests were performed using isolated colonies from BHI agar plates and *L. monocytogenes* 10403S as a control. For the catalase test, 3% hydrogen peroxide (Medique Products, Fort Myers, USA) was added and observed for the formation of gas bubbles. For the oxidase test, an oxidase test strip (OxiStrips; Hardy Diagnostics, Santa Maria, CA) were used.

### Hemolysis

Hemolysis was evaluated by stab inoculating sheep blood agar plates (SBA; commercially prepared; Hardy Diagnostics, Santa Maria, CA) from BHI agar plates. *Listeria monocytogenes* 10403S, *Listeria monocytogenes* ATCC 19115, *Listeria ivanovii* ATCC 19119, and *Listeria seeligeri* ATCC 35967 were included as positive controls and *Listeria innocua* ATCC 33090 as a negative control. SBA plates were incubated aerobically at 35°C and checked at 24 and 48 h.

### Nitrate reduction

Nitrate reduction was evaluated by inoculating nitrate broth with Durham tubes (commercially prepared; COMPANY, LOCATION) with several colonies from BHI agar and incubating at 35°C for up to 7 d. Reduction of nitrate to nitrite was evaluated daily by adding NIT1 and NIT2 reagents (bioMérieux, Marcy-l’Étoile, France) to 1 mL aliquots. If negative, zinc powder (bioMérieux, Marcy-l’Étoile, France) was added to confirm the presence of nitrate. Additionally, tubes were observed for the production of gas, which would indicate further reduction to gaseous nitrogen products. *L. booriae* FSL A5-0281^T^ and *L. monocytogenes* 10403S were included as positive and negative controls, respectively.

### Biochemical test kits

API *Listeria*, API 20 E and API 50 CH test kits (bioMérieux, Marcy-l’Étoile, France) were all performed following the manufacturer’s instructions. For the API 50 CH strips, the *Bacillus* methods in the instructions and API 50 CHB/E medium were used. *L. monocytogenes* 10403S and *L. innocua* ATCC 33090 were used as controls for all three, in duplicate. Inoculum for the strips was prepared by suspending colonies from BHI agar in the appropriate suspension medium for each. API *Listeria* and API 20 E strips were incubated at 35°C for 24 h and then interpreted. API 50 CH test strips were incubated at 30°C and interpreted at 24 and 48 h.

## RESULTS/DISCUSSION

Since 2010, there have been multiple new species added to the *Listeria* genus, many originally isolated from natural environments^16^. This paper describes the genotypic and phenotypic characterization of two new *Listeria* isolates obtained from soil samples collected in the Great Smoky Mountains National Park along the North Carolina-Tennessee border^17^. Evaluation of genotypic and phenotypic characteristics of “L. swaminathanii” strains will aid in the characterization of this novel species and contribute to our knowledge of the diversity of *Listeria* spp. Here, we describe the newly isolated strains, UTK C1-0015 and UTK C1-0024, and compare with the “L. swaminathanii” type strain (FSL L7-0020^T^) and other *Listeria* spp.

Both genomes were able to be assembled into complete closed genomes (contiguous sequences that comprise the entire genome). The genome of UTK C1-0015 consists of a 2.78 Mb chromosome and 55 Kb plasmid (total genome length of 2.84 Mb) with a G+C content of 38.7%; UTK C1-0024 consists of a 2.95 Mb chromosome with a G+C content of 38.6% (**Table 1**), which is consistent with FSL L7-0020^T^. Of the validly published type strains, the two isolates showed highest similarity to *L. marthii* (94.0-94.1%) (**Figure 1**); however, they were most closely related to “L. swaminathanii” FSL L7-0020^T^, with 98.7-98.8% ANI, indicating that they belong to the same species. Examination of the chromosomal alignment of the two isolates and the type strain shows that, overall, there is a high level of conservation across the entire chromosome, with no large rearrangements or deletions (**Figure 2**). However, there are some loci throughout that are present or absent in only one of the isolates.

**Table 1.**
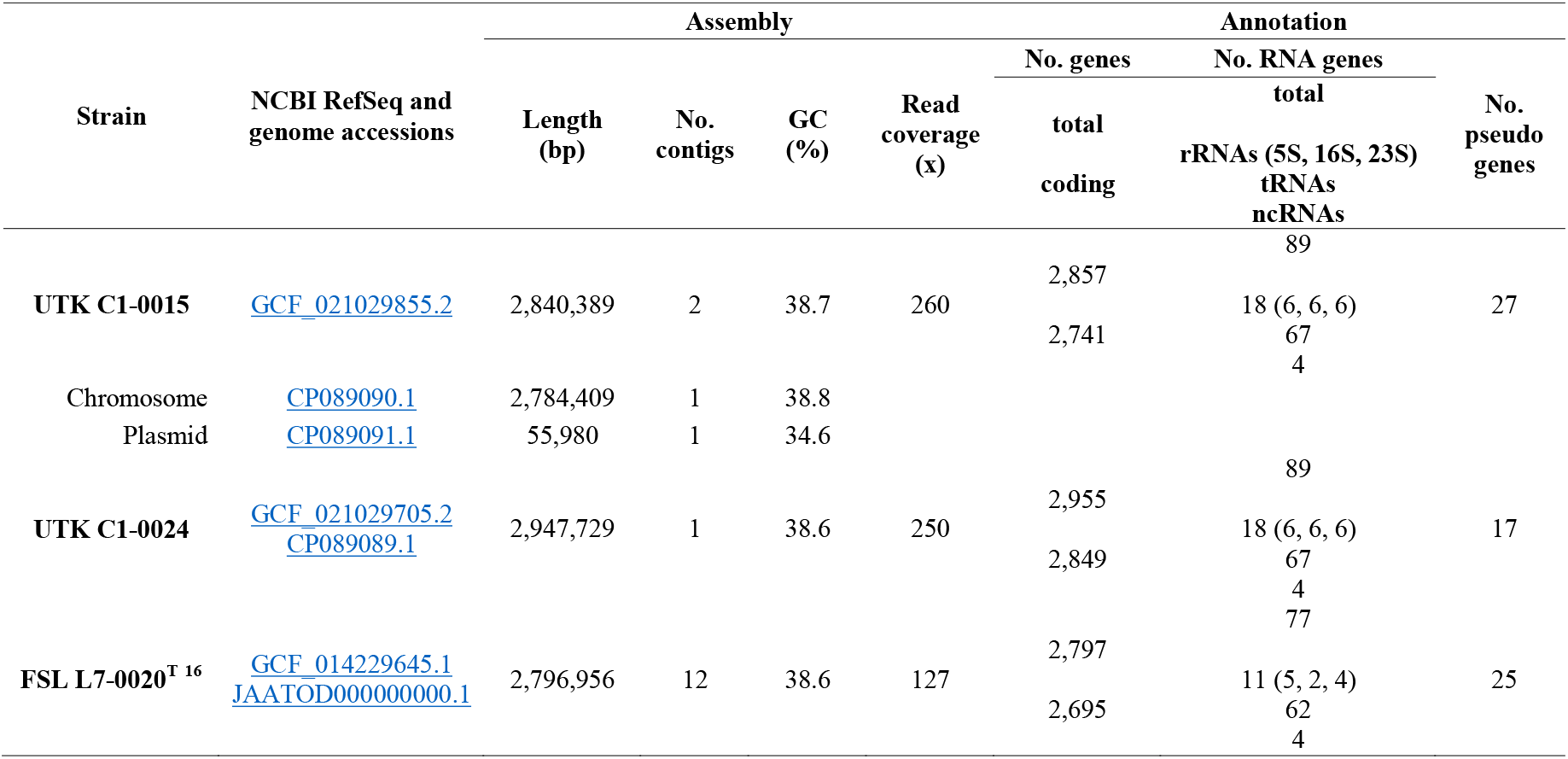
Genome Statistics Genome statistics for the two “L. swaminathanii” strain hybrid assemblies described here and for the recently described type strain^16^.

**Figure 1.**
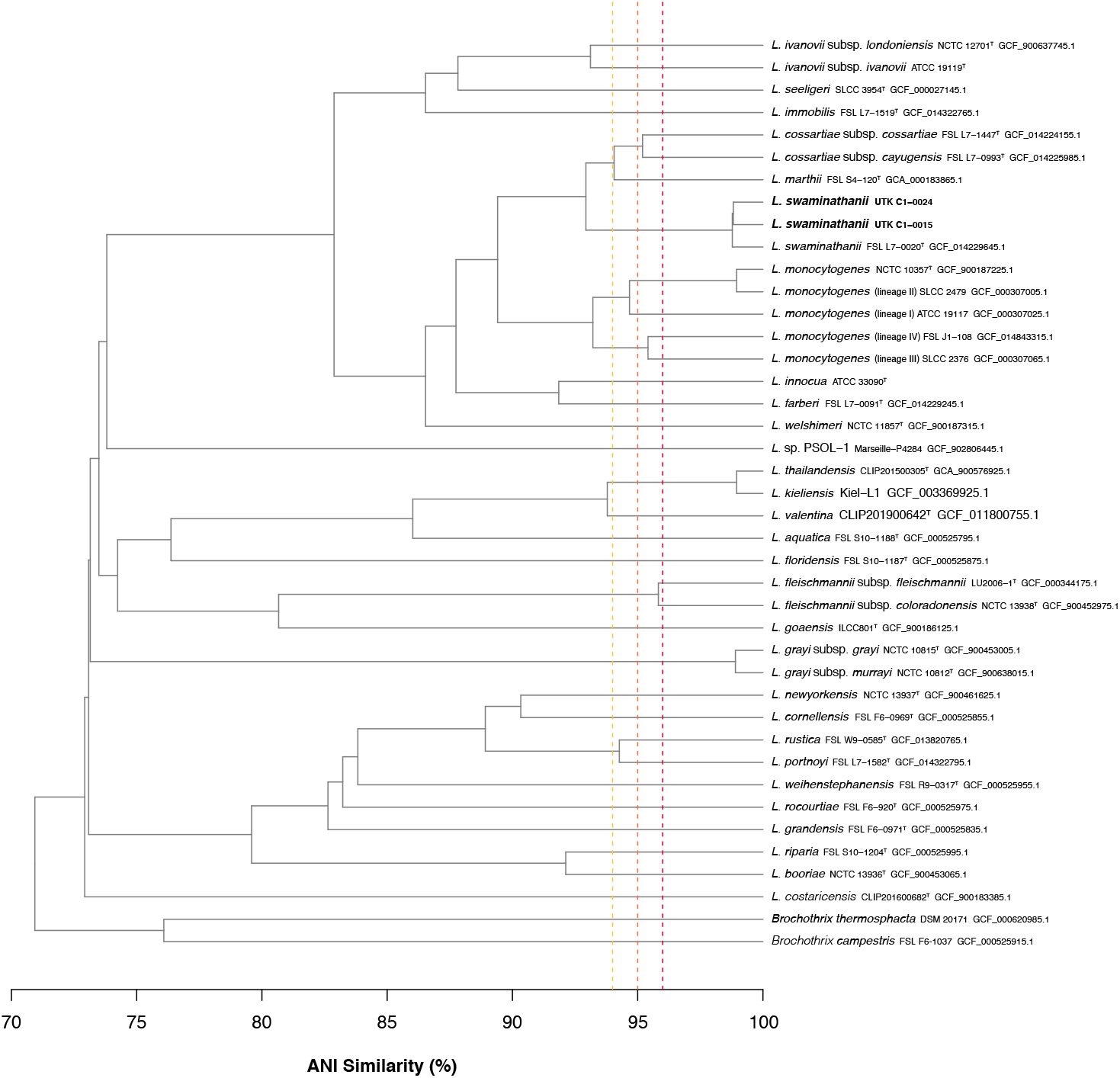
ANI Similarity Dendrogram Average nucleotide Identity (ANI) dendrogram of the recently isolated “L. swaminathanii” strains (bold), along with all described *Listeria* spp. type strains and representative from each of the *L. monocytogenes* lineages (indicated in parentheses). Horizontal distance represents ANI similarity (%) and vertical dashed lines indicate ANI values of 96 (yellow), 95 (orange), and 94% (red).

**Figure 2.**
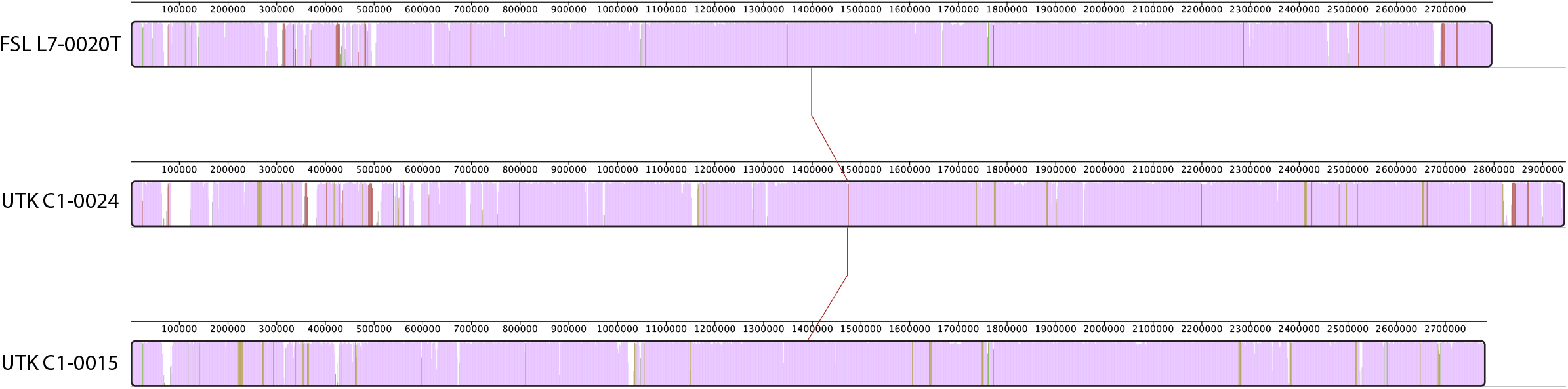
Chromosomal alignment of FSL L7-0020^T^, UTK C1-0024, and UTK C1-0015. Alignment shows three horizontal panels, one per strain. The colored portions inside each panel represents sequence similarity, with height corresponding to average conservation at that location. Regions that are conserved among all genomes are purple. Regions that are conserved among only two of the genomes are red (FSL L7-0020T and UTK C1-0024), green (FSL L7-0020^T^ and UTK C1-0015), or yellow (UTK C1-0024 and UTK C1-0015). Regions without coloring were not aligned and likely contain loci that are present in only a single genome.

Both genomes contained the following antibiotic resistance genes: *fosX, lin, norB*, and *sul*. Virulence-associated genes involved with adherence (*dltA, fbpA, lap, lapB, pdeE*), bile-resistance (*bsh, mdrM*), immune modulation (*lntA*), intracellular survival (*lplA1, oppA, pdeE, prsA2, purQ, svpA*), invasion (*iap, lpeA, pdeE*), peptidoglycan modification (*oatA, pdgA*), regulation of transcription and translation (*agrAC, cheAY, codY, fur, lisKR, stp, virRS*), surface protein anchoring (*lgt*, *lspA, srtAB*), and teichoic acid biosynthesis (*gltB, gtcA*) were identified in both genomes, along with internalins *inlGHJK, inlC2*, and *inlD* (**Supplementary Table S1**). Genes associated with *Listeria* pathogenicity island LIPI-3 (*llsABDGPXY*) were only found in UTK C1-0024 (**Supplementary Figure S1**), as well as *gltA* (teichoic acid biosynthesis). The internalin genes *inlA* and *inlB* and genes associated with *Listeria* pathogenicity islands LIPI-1, LIPI-2, or LIPI-4 were not detected in either.

A 56 Kb plasmid was identified in UTK C1-0015. The plasmid has an Illumina read depth of 2.2x the overall median depth, indicating a copy number of two. The plasmid found in UTK C1-0015 shows a high similarity (86.03% nucleotide identity) to pLMIV from *L. monocytogenes* strain FSL J1-0208^46,47^. However, pLMIV is approximately 21 Kb longer than the plasmid found in UTK C1-0015; this is due to the presence of a region encoding four complete internalins and one internalin-like protein in pLMIV, this region is absent in the plasmid in UTK C1-0015 (**Figure 3**). Both plasmids are also similar to the plasmid in *L. monocytogenes* FSL J1-0158. Both FSL J1-0208 and FSL J1-0158 were originally isolated from clinical caprine sources^46^. Most genes in the plasmid found in UTK C1-0015 seem to encode proteins predicted to be involved in plasmid maintenance and conjugation^46^, with only a few putative cargo genes, most which are of unknown function and one encoding a DNA-methyltransferase.

**Figure 3.**
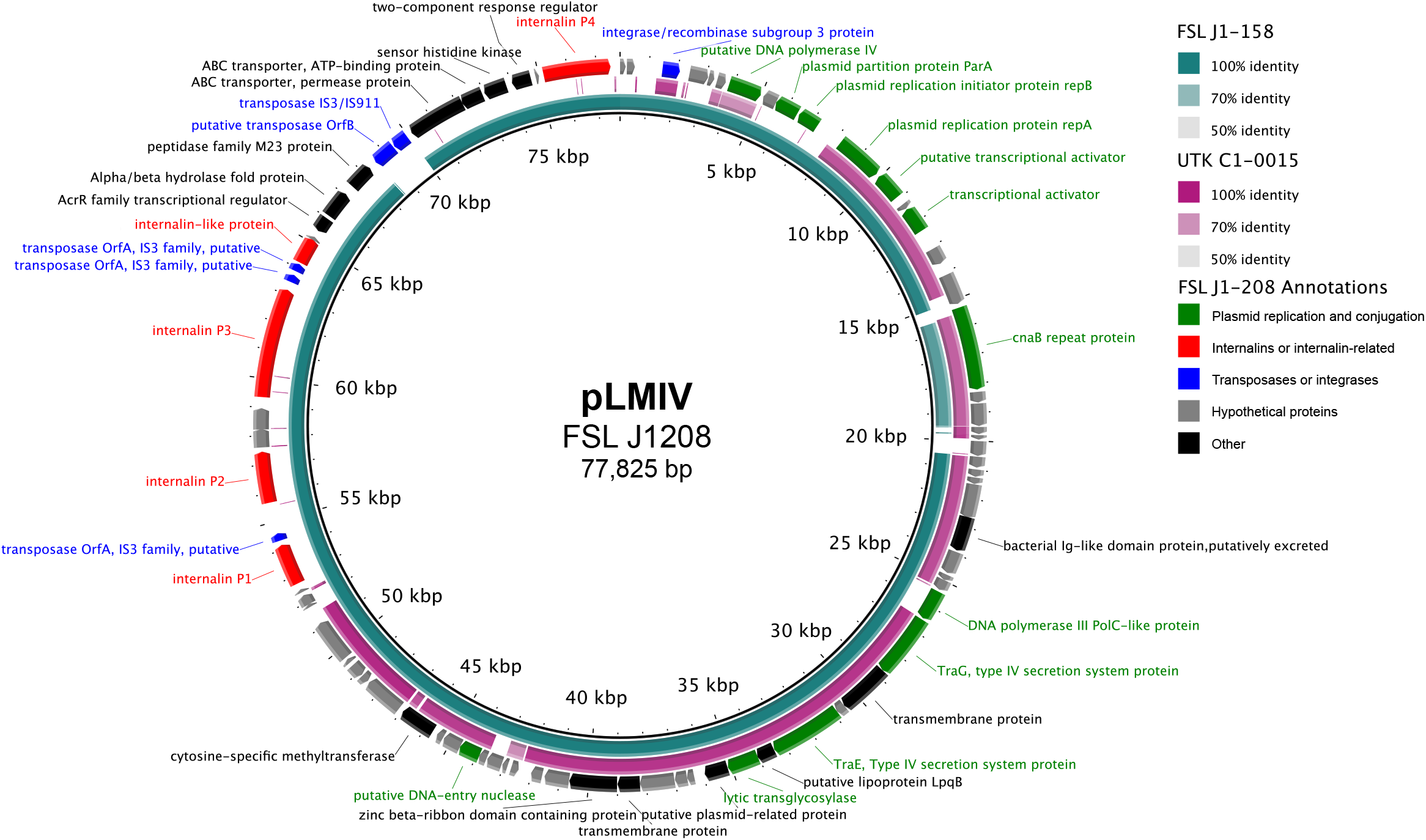
Comparison of plasmid found in UTK C1-0015 to plasmids from FSL J1-020 and FSL J1-158. Comparison of the plasmids found in UTK C1-0015, FSL J1-020, and FSL J1-158, using pLMIV from J1-208 as the reference. The innermost black ring represents pLMIV. The middle rings represent FSL J1-158 (teal) and UTK C1-0015 (purple), with BLAST identity indicated by shading (see legend). The outermost ring contains gene annotations from pLMIV that are colored by functional category: green (plasmid replication and conjugation), red (internalins or internalin-related), blue (transposases or integrases), gray (hypothetical proteins), and black (other).

PHASTER and PhageBoost were used to predict prophage sequences in the genomes. The genome UTK C1-0024 was also predicted to house a prophage integrated near a tRNA-Lys gene. Blastn results show the prophage from UTK C1-0024 has an 88.58% identity to *Listeria* phage A500 with 60% coverage. Prophages and other mobile genetic elements can contribute to genome diversity and have been used to distinguish epidemic clones of *L. monocytogenes*^48–50^. Strain UTK C1-0015 was predicted to house a partial monocin locus of eight open readings frames^51,52^; structural genes such as those that code for the tail tape measure protein or tail fibers were absent from the locus. The monocin locus from strain UTK C1-0015 shares a 99.405% identity to the monocin locus from FSL L7-0020^T^ (GCF_014229645.1). The UTK C1-0024 genome was predicted to house the full monocin locus of 18 open readings frames, similar to the monocin in *L. monocytogenes* strain 10403S **(Figure 4)**. Blastp queries using the monocin locus from UTK C1-0015 and UTK C1-0024 return hits to *L. marthii, L. cossartiae, L. innocua, L. farberi*, and *L. monocytogenes* strains with 100% coverage and >89.90% identity, suggesting this is fairly dispersed across the *sensu stricto* clade of *Listeria*. Monocins are bacteriocins produced by the host that may be significant in establishing dominant strains in ecological niches, as they target closely related species, but remain inactive against the producing strain^53^.

**Figure 4.**
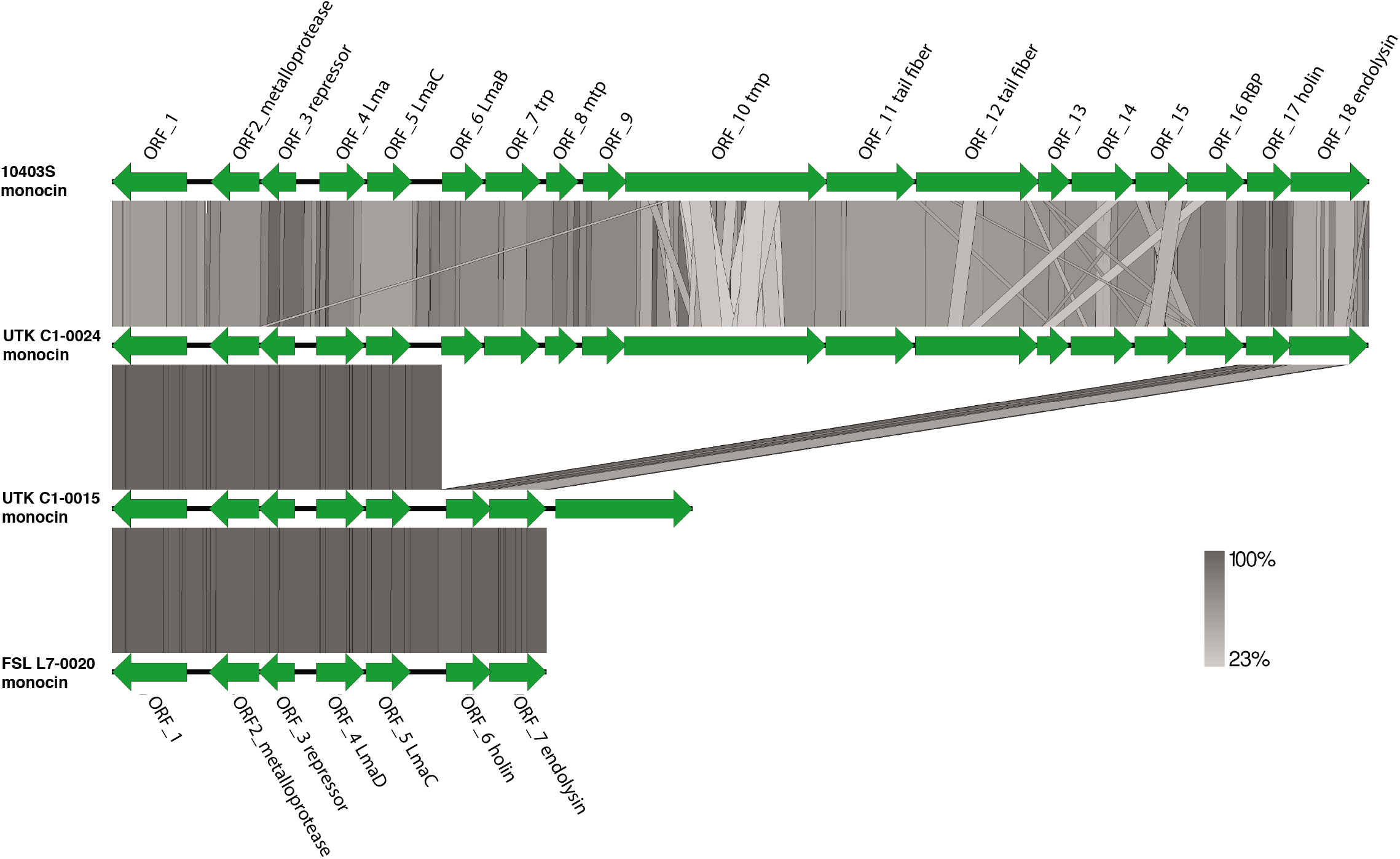
Nucleotide similarity of monocin regions BLAST comparisons of monocin regions from *L. monocytogenes* 10403S, UTK C1-0015, UTK C1-0024, and the “L. swaminathanii” type strain FSL L7-0020^T^. Genes are represented by green arrows. The shaded regions represent nucleotide similarity (see scale at bottom right).

*Listeria* spp. grow at a wide range of temperatures from 0-45°C^7,8,14,16^ and can survive at temperatures below freezing (−7°C)^54^. In the current study, we performed growth assessments at 4, 7, 22, 30, 37, and 41°C. These temperatures were chosen to encompass the known growth temperature range, with 4 and 7°C specifically included because some species are unable to grow well at low temperatures (<7°C)^4^. Strains UTK C1-0015 and UTK C1-0024 exhibited growth at all temperatures tested (**Supplementary Table S2**), except for UTK C1-0024 at 41°C. After 24 h of incubation, UTK C1-0015 and UTK C1-0024 showed optimal growth at 30°C (9.2 and 9.4 log_10_ CFU/mL), followed by at 37°C (8.9 and 9.0 log_10_ CFU/mL). At 41°C, no growth was observed for UTK C1-0024 at 41°C, which is dissimilar to both UTK C1-0015 and FSL L7-0020^T^. At 4°C, the concentration increases of UTK C1-0015 and UTK C1-0024 were higher than the increases seen in FSL L7-0020^T^.

*Listeria* spp. are Gram-positive rods^7^; this was confirmed for UTK C1-0015 and UTK C1-0024. Both isolates were observed to grow under aerobic and anaerobic conditions at 30°C after 24h; this is another expected result, as *Listeria* spp. are facultative aerobes^7^. Both strains were oxidase negative (**Supplementary Table S3**), as expected^7^, indicating a lack of cytochrome c oxidase. Additionally, both were catalase positive, indicating they produce the catalase enzyme that converts hydrogen peroxide into oxygen gas and water; however, FSL L7-0020^T^ is catalase negative^16^, a phenotype that has only been described in one other *Listeria* spp. (*L. costaricensis*)^55^. When kat gene from the reference, two isolates, and the type strain are aligned, there are nucleotide differences at 158 positions. 16 of the nucleotide differences differ between the type strain and one or both of the isolates. Four of those result in amino acid differences, with two between the type strain and both isolates. At amino acid position 72, the type strain has glutamic acid (polar, acidic) and the two isolates have lysine (polar, basic), a radical substitution. At amino acid position 92, the type strain has histidine and the other two arginine (both polar, basic), a conservative substitution. These amino acid differences may have an effect on the structure and function of the resulting protein, leading to the catalase-negative phenotype of FSL L7-0020^T^.

On MOX agar, UTK C1-0015 and UTK C1-0024 colonies were typical for *Listeria* spp.: gray to black colonies with sunken centers and black halos, indicating esculin hydrolysis. On *Listeria* CHROMagar, UTK C1-0015 and UTK C1-0024 were typical for *Listeria* spp.: blue colonies (indicating β-glucosiadase enzyme activity), but lacking opaque white halos typical for *L. monocytogenes* and *L. ivanovii* (indicating no phosphoatidylinositol-specific phospholipase C [PI-PLC] activity) (**Supplementary Table S3**).

API test kits were used to characterize metabolic function of UTK C1-0015 and UTK C1-0024. The *Listeria* API kit is designed for species-level identification *Listeria* spp. based on enzymatic tests and sugar fermentations. For this test, both strains generated a code of 6110 (**Supplementary Table S3**), consistent with FSL L7-0020^T 16^ and indicates an 80% (t-value of 0.62) ID to *L. monocytogenes* according to the APIweb database. The control strains, *L. monocytogenes* 10403S and *L. innocua* ATCC 33090, generated the expected codes of 6510 and 7510, respectively.

The API 20 E kit is designed for identification of Enterobacteriaceae and other non-fastidious Gram-negative rods; however, this kit contains tests that can be used for genus-level identification of *Listeria* spp. and has been used previously in the characterization of novel *Listeria* spp.^10,16^. For this test, UTK C1-0015 and UTK C1-0024 were positive for acetoin production (Voges Proskauer) and D-glucose and amygdalin fermentation, which is consistent with *L. monocytogenes* 10403S, *L. innocua* ATCC 33090, and FSL L7-0020^T 16^ (**Supplementary Table S3**). UTK C1-0015 and UTK C1-0024 were negative for all other tests, including indole, urease, and H_2_S production.^16^ All API 20 E results were consistent with FSL L7-0020^T 16^. Nitrogen reduction was evaluated using both the API 20E kits and nitrogen broth; both strains were negative.

The API 50 CH kit is designed for the study of carbohydrate and carbohydrate-derivative metabolism and API 50 CHB/E medium is designed for use with *Bacillus* and related genera, *Enterobacteriaceae*, and *Vibrionaceae*. Results for this test were consistent between UTK C1-0015, UTK C1-0024 and FSL L7-0020^T^ (**Supplementary Table S3**), with four differences. UTK C1-0015 yielded a negative result for D-lactose, a result that differs from UTK C1-0024, FSL L7-0020^T^, and most *sensu stricto Listeria* species^16^. Both strains tested negative for glycerol and starch (amidon); this differed from the type strain^16^, which is positive for both. UTK C1-0024 was positive for D-trehalose fermentation, while UTK C1-0015 and the type strain were negative. Examination of the genomes shows that a locus containing three genes associated with trehalose fermentation (*treR, treC*, and *treP*) is present in UTK C1-0024, but absent in the two other genomes. In *L. monocytogenes*, trehalose has been shown to increase biofilm formation^56^. The API 50CH test is a qualitative test and interpretation of results can vary, which is one major limitation of qualitative tests.

The complete lysis of red blood cells, β hemolysis, is associated with pathogenicity in *Listeria* spp.^7^ On SBA, UTK C1-0015 and UTK C1-0024 were non-hemolytic, which is consistent with the non-hemolytic FSL L7-0020^T 16^ and the negative control *L. innocua* ATCC 33090. β hemolysis is typically only observed in *L. monocytogenes, L. ivanovii*, and *L. seeligeri*.^7,45^

When observed microscopically, both UTK C1-0015 and UTK C1-0024 appeared motile at 25°C and nonmotile at 37°C (**Supplementary Table S3**). Motility at 25°C was confirmed with MTM tubes; both strains were clearly motile after 5 d of incubation as evidenced by an umbrella-shaped growth pattern, characteristic of motile *Listeria* spp. These results were consistent with FSL L7-0020^T 16^ and other *sensu stricto* species, with the exception of *L. immobilis* (non-motile at 25°C^10^). In *L. monocytogenes*, motility genes like flagellin are expressed at lower temperatures like 25°C, but become restricted at 37°C^57^.

### Conclusions

In this study, we described two strains isolated from soil samples collected in the GSMNP, which belong to the recently proposed novel species “L. swaminathanii”. By the addition of two additional strains to this species (bringing the total number described to three), the diversity of this species can be further evaluated and the characteristics of these two strains can be compared to those of the type strain FSL L7-0020^T^. Additionally, we were able to provide complete closed genomes for both strains, including the plasmid found in UTK C1-0015, and further characterize genomic features. As the two strains described in this study were also isolated from the GSMNP, they are subject to the same restrictions as the proposed “L. swaminathanii” type strain.

## ADDITIONAL INFORMATION

### Data Availability

Raw sequencing reads and genome assemblies are available on NCBI under BioProject PRJNA760531.

## Acknowledgements

We thank Martin Wiedmann (Cornell University) for providing the *Listeria booriae* strain FSL A5-0281.

## Author contributions

Conceptualization: LKH, CRC, HCdB, TGD; data curation: LKH, HKC, CS, TLP; formal analysis: LKH, TLP; funding acquisition: TGD; investigation: LKH, HKC, CS, MLC, DWB, TLP, YS; methodology: LKH, HKC, HCdB, TGD; project administration: LKH, DWB, TGD; resources: TGD; supervision: LKH, TGD; visualization: LKH, HKC, CS, TLP; writing (original draft): LKH, HKC, CS, TLP; writing (review & editing): LKH, HKC, CS, MLC, DWB, TLP, YS, CRC, HCdB, TGD.

