## Supplemental Figure S1 for "Phenotypic characterization and analysis of complete genomes of two distinct strains of the proposed species “L. swaminathanii”"

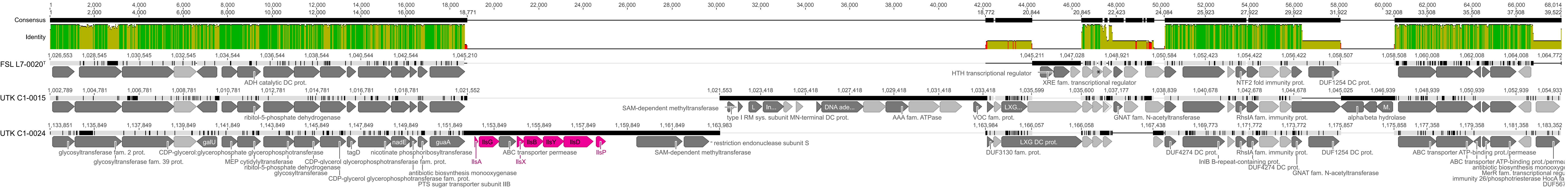


**Supplementary Figure S1.** Listeria pathogenicity island 3 (LIPI-3) in UTK C1-0024

Region of the whole-genome alignment (Figure 1) that contains LIPI-3 (CDS in pink). CDS annotated as hypothetical proteins are light gray and all others are dark gray.
